# Transformation of Independent Oscillatory Inputs into Temporally Precise Rate Codes

**DOI:** 10.1101/054163

**Authors:** David Tingley, Andrew A. Alexander, Laleh K. Quinn, Andrea A. Chiba, Douglas Nitz

## Abstract

Complex behaviors demand temporal coordination among functionally distinct brain regions. The basal forebrain’s afferent and efferent structure suggests a capacity for mediating such coordination. During performance of a selective attention task, synaptic activity in this region was dominated by four amplitude-independent oscillations temporally organized by the phase of the slowest, a theta rhythm. Further, oscillatory amplitudes were precisely organized by task epoch and a robust input/output transform, from synchronous synaptic activity to spiking rates of basal forebrain neurons, was identified. For many neurons, spiking was temporally organized as phase precessing sequences against theta band field potential oscillations. Remarkably, theta phase precession advanced in parallel to task progression, rather than absolute spatial location or time. Together, the findings reveal a process by which associative brain regions can integrate independent oscillatory inputs and transform them into sequence-specific, rate-coded outputs that are adaptive to the pace with which organisms interact with their environment.

---

Encoding and passage of information via rhythmic electrical activity patterns in the brain provides an efficient means by which to transmit information, one complementary to encoding of information in spike rates. The brain appears to take great advantage of neural oscillations in information transfer and such mechanisms may be dysfunctional in neurological disorders such as schizophrenia and autism^1^,^2^,^3^,^4^,^5^.

Oscillations in action potential discharge in connected brain regions are often highly coherent across narrow frequency-ranges in relation to specific cognitive processes such as attention^6^,^7^. Yet, prominent neural oscillations occur at a number of different frequencies^8^,^9^,^10^,^11^ and it remains to be determined how they are organized and decoded within a single recipient system. In particular, for ‘associative’ brain regions integrating a wide array of inputs, it is unclear whether and in what form coordination of different oscillatory inputs is achieved. Equally, it is unclear how oscillatory input amplitudes may be translated into other forms of neural information, such as rate-coded sequences of cell assemblies^12^.

Valuable clues as to how such processes could occur may be found in telecommunications where communication across a network of devices must be accomplished along bandwidth-limited pathways^13^,^14^. Here, coordination through multiplexing of different sources of information in distinct oscillatory frequencies provides a powerful means by which to transmit independent information sources^15^. Yet, critical questions remain as to whether specific information transmission models such as time-division multiplexing and frequency-division multiplexing are applicable to brain networks and whether robust evidence for their utilization is found during performance of complex tasks.

With regard to these questions, the mammalian basal forebrain (BF) represents an excellent test case with which to consider how oscillatory inputs may be integrated and transformed into sequenced ensemble firing rate patterns. First, the BF receives afferents from a remarkably wide array of cortical and sub-cortical brain regions^16^, many of which exhibit oscillatory spiking activity across a wide range of frequency bands^17^,^18^,^19^,^20^,^21^,^22^. Second, individual BF neurons exhibit stimulus-induced oscillatory spiking in vitro^23^. Thus, at least some BF synaptic inputs, whether intrinsic or extrinsic in source, are likely to be organized as oscillations and to therefore be observed in recordings of local field potentials (LFPs). Third, BF ensembles generate robust sequences of rate-coded output that strongly correlate with specific epochs of tasks requiring selective attention, stimulus encoding, decision-making, and outcome evaluation^24^. Fourth, such BF neuron ensembles operate as cell assemblies in vivo^25^, indicating the capacity for integration of oscillations at both the cellular and network levels. Finally, BF outputs reach a wide array of targets, implying a role in coordinating activity patterns among widely separated brain regions based on the integrated activity from an equally wide array of afferents^26^,^27^,^28^,^29^.

Consistent with this premise, examination of BF local field potentials revealed prominent oscillations in four distinct frequency bands (Fig. 1A). The higher frequency oscillations, beta (20-35 Hz), gamma (45-65 Hz), and hi-gamma (80-150 Hz) often occurred as transient events (Fig. 1B) and were independent of each other in their power fluctuations (Fig. 1C). The observed features are hallmarks of frequency-division models for multiplexing where different information sources are organized at different frequencies and with independence in variation of their amplitudes. Beta, gamma, and hi-gamma transients were also temporally organized across the phases of individual theta waves (4-9 Hz; Fig. 1D and Fig. S1). The observed temporal framework of phase/amplitude relationships suggests that the BF theta rhythm is responsible for sequencing higher frequency transients through time. This latter feature is a hallmark of time-division multiplexing^15^ where different information sources reach their target according to an ordered sequence.

**Figure 1.**
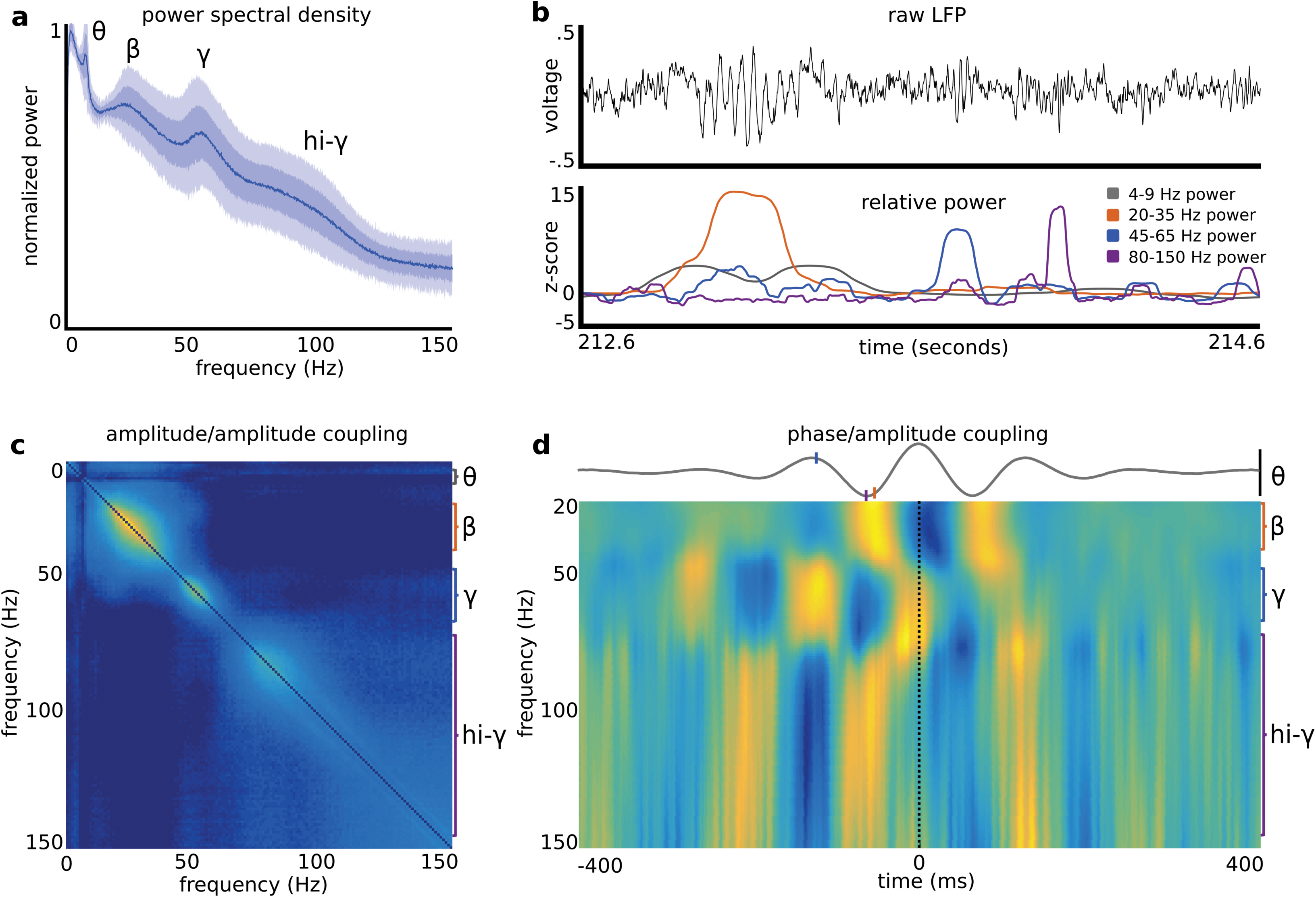
Beta, gamma, and hi-gamma transients occur within a temporal framework that is referenced to theta. **(A)** Average power spectral density plots for 36 recordings from four animals. Dark and light transparent shading represent one and two standard deviations respectively. Plots were calculated using local field potential data from entire recording sessions (~25-40 minutes). Power spectral density plots were also remarkably consistent across recording days in single individuals (Fig. S1A). **(B)** Example 2-second raw LFP trace (black), and z-scored power at theta (grey; 4-9 Hz), beta (orange; 20-35 Hz), gamma (blue; 45-65 Hz), and hi-gamma (magenta; 80-150 Hz) frequency bands. **(C)** Average power/power correlations for all animals across all frequencies, showing that there are multiple frequency bands that are correlated within a given range but independent from other frequency ranges. Color axis (0-0.5). **(D)** Phase/amplitude coupling to theta (4-9 Hz) frequency. The average theta wave, aligned by the peaks, across all recordings is plotted as the grey line. The colormap represents the average wavelet transform relative to the peak of theta. Beta (20-35 Hz) and high gamma (80-150 Hz) frequencies increase in amplitude during troughs in theta, while gamma power (45-60 Hz) increases at the peak of theta oscillations. Orange, blue, and magenta ticks on the average theta wave indicate maximum power for beta, gamma, and hi-gamma frequency bands respectively. Black vertical bar for theta wave (−2 to +2 microvolts). Values at each frequency are normalized to the highest value at the same frequency. Color axis (0-1).

Notably, time-division and frequency-division models for decoding and/or integration of separate information sources are not mutually-exclusive. In addition, coherence of spiking activity to the phase of ongoing oscillatory inputs is consistent with both forms of multiplexing. Accordingly, in the present sample, spiking activity for a large sub-population of BF neurons was robustly phase locked to the observed LFP frequency bands (Fig. 2). Across the population, neurons varied widely with respect to their spiking coherence to the four major LFP oscillatory frequency bands with some neurons showing coherence to more than one frequency and some to none. Together, then, it is clear that both BF oscillatory inputs and BF spiking are organized in a manner consistent with at least two, notably non-mutually-exclusive, models for integration of multiplexed information.

**Figure 2.**
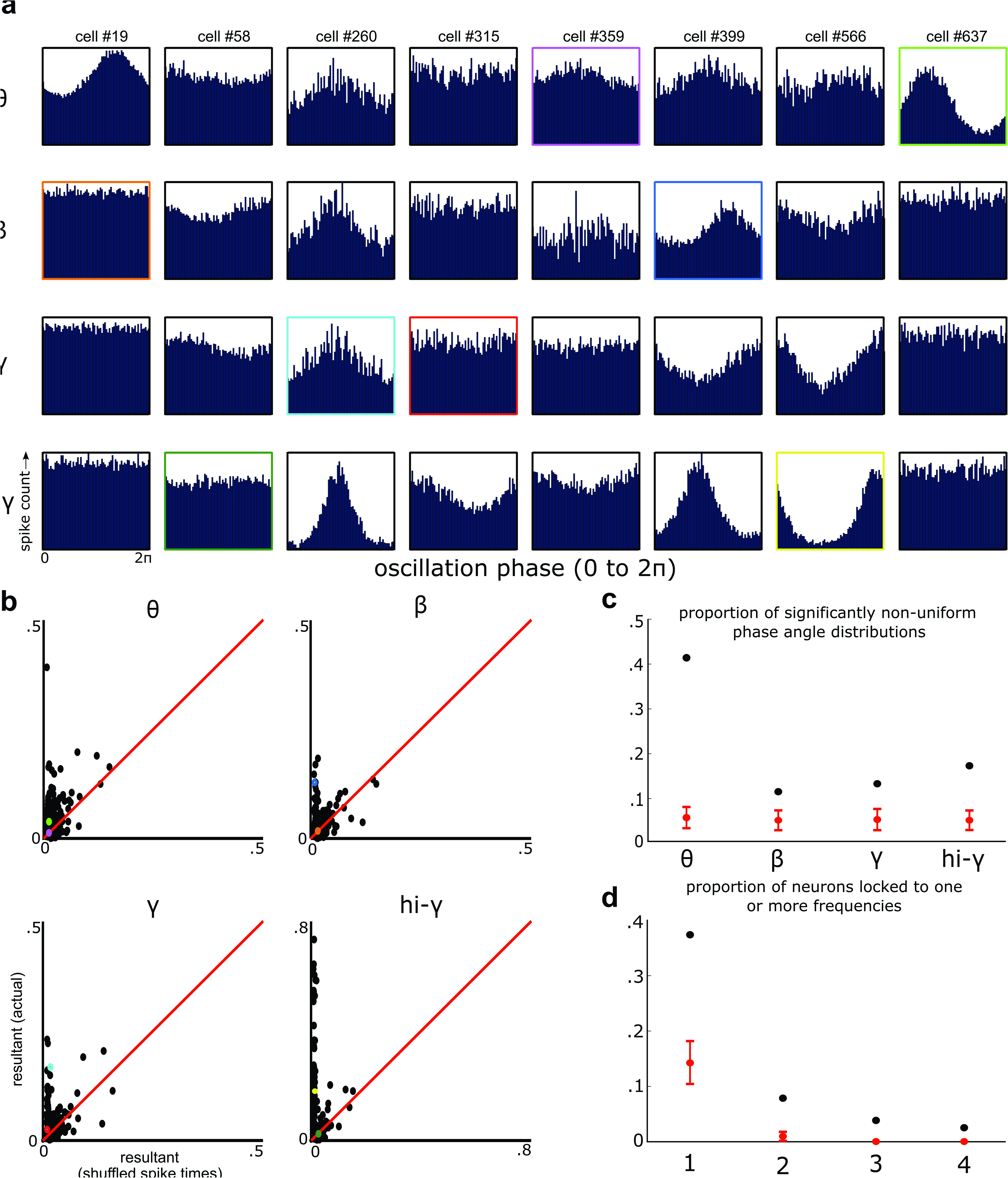
Individual BF neurons phase lock to specific frequency bands. **(A)** For each example neuron (columns), there are four plots (rows) that show the spike time histograms relative to the phase of theta, beta, gamma, and hi-gamma oscillations. Colored boxes map onto the colored dots in Fig. 2B. **(B)** For each neuron, resultant vectors were calculated for the distribution of phase angles at spike times and for randomized spike times (avg. of 100 iterations). Scatter plots depict the actual resultant vector (y-axis) and the average resultant vector with randomized spike times (x-axis) for each neuron. Red lines indicate a slope of 1 and colored dots map onto the colored outlines in Fig. 2A. Light green, blue, teal, and yellow dots correspond to strongly phase locked neurons for theta, beta, gamma, and hi-gamma respectively; while magenta, orange, red, and dark green dots correspond to neurons with weak or no modulation for theta, beta, gamma, and hi-gamma respectively. **(C)** For each neuron, Rayleighs test for non-uniformity was then used to determine whether spike times were uniformly distributed, or locked to particular phases of theta, beta, gamma, or hi-gamma oscillations. The proportion of neurons with p-values < .05 are represented as the black dots for each frequency band. In red, is the mean proportion of neurons with p-values < .05 when spike times are randomly shuffled (100 iterations; error bars are +/- 3 standard deviations). **(D)** The proportion of neurons with significantly non-uniform spike phase distributions for one, two, three, or four of the observed frequency bands are shown as the black dots. In red, is the proportion of neurons with p-values < .05 for one, two, three, or four frequency bands when spike times are randomly shuffled (100 iterations; error bars are +/- 3 standard deviations).

The foregoing data suggest a capacity for BF networks to efficiently integrate independently-varying sources of input to, in turn, produce spiking output consistent with coordination of responsiveness in efferent targets such as cortex. However, a next critical step was to determine whether and how such a process is actually utilized during performance of a task involving a behavioral series associated with variation in sensory stimuli, required actions, and cognitive demands. Therefore, LFP and spiking activity relationships during a multi-epoch selective attention task were analyzed in order to determine: 1) whether LFP oscillations themselves are organized relative to the cognitive demands associated with different epochs of task performance; and 2) whether these signals may be integrated or transformed into other coding regimes as part of BF spiking output (Fig. 3A).

**Figure 3.**
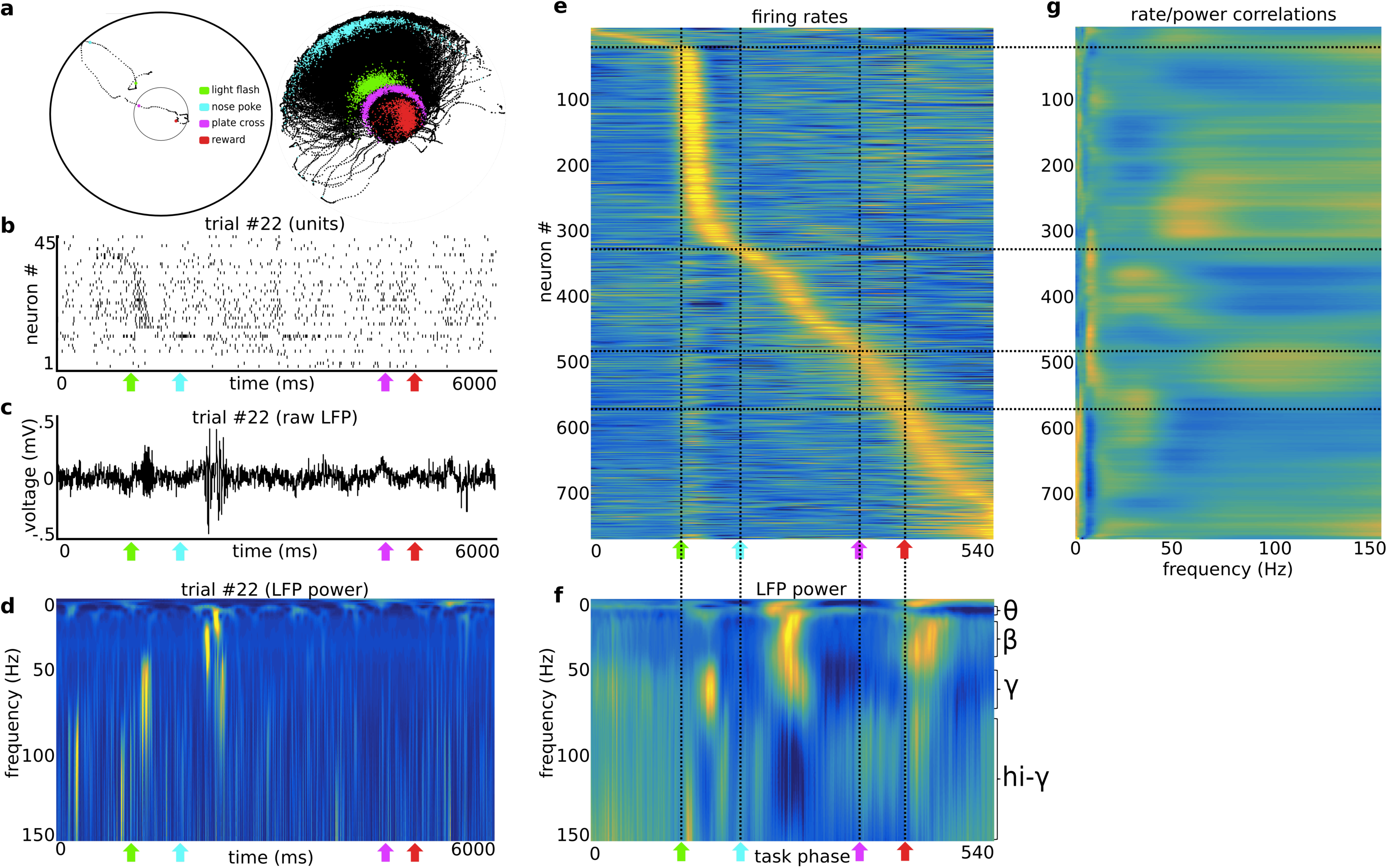
Neuronal firing is correlated with the oscillation amplitude of specific frequency bands. **(A)** Example tracking data from a single trial (left) and 24 recordings (right). Head location was detected via two head-mounted LED’s (black dots). Green, cyan, magenta, and red dots indicate the head location during the light flash, nose-poke, plate cross, and reward location respectively. **(B)** Example spike trains from 45 simultaneously recorded BF neurons across a single trial. **(C)** Raw LFP trace from the same example trial as in 2B. **(D)** Wavelet transform of LFP trace in 2C. Note the gamma and beta transients after the light flash and nosepoke respectively (color axis 0-1 - maximum normalization performed individually for each frequency bin). **(E)** Mean firing rates for 780 BF neurons. Each row is the average firing rate (~70 trials) for a single neuron; the population is sorted by the point of maximum firing during the selective attention task (color axis 0-1 with each maximum normalization performed individually for each row/neuron). For further analysis of task-epoch-specific firing, see Tingley et al. 2014. **(F)** Average wavelet transform of local field potentials recorded during the selective attention task. Changes in local field potential power during the task can be observed across all four frequencies; theta (4-9 Hz), beta (20-35 Hz), gamma (45-60), and high gamma (80-150 Hz). All frequencies (rows) are individually maximum-normalized (color axis is 0-1) to visualize changes across the spectrum of frequencies present. **(G)** Neurons with specific firing patterns, relative to the selective attention task (3E), correlate with specific frequency bands in the local field potential (3F). For each trial, the cross-epoch firing rate vector for each neuron is correlated (Pearsons product-moment correlation) with the wavelet transform of the local field potential across the same epochs. These correlations are then averaged across trials, generating a matrix where the y-axis is neuron number (1-780) and the x-axis is local field potential frequency (1-150 Hz). The rows of this correlation matrix are then sorted to match the order of 3E, maximum normalized, and smoothed with the nearest 30 neurons (moving window along the y-axis; color axis is between 0 and 1). For comparison with correlation values expected by chance, see Fig. S3. Black vertical lines (across 3E and 3F) mark the light flash, nosepoke, plate cross, and reward; while the black horizontal lines (across 3E and 3G) mark the neurons with peak firing rates closest to these behaviorally defined events.

Animals completed a selective attention task where they identified transient light flashes on the perimeter of a circular environment with a nose-poke at the light location (Fig. 3A; Tingley et al. 2014). Across all trials associated with success in light source identification and reward delivery (~70 per recording), the spiking activity of single neurons followed stereotyped patterning relative to task epoch (Fig. 3B; see also Tingley et al. 2014). Prominent oscillatory LFP transients occurred during individual trials (Fig. 3C-D), and with high trial-to-trial and animal-to-animal reliability in their association to specific task phases (Fig. 3F, Fig. S2). Thus, the independent oscillatory LFP components identified in figure 1 are indeed relevant to the variations in experience and cognitive processing associated with performance of this task. Furthermore, the finding that both neural spiking and LFP oscillations follow stereotyped task-epoch-specific patterns, suggests that a direct coupling of oscillation amplitude and firing rate might characterize the transformation of multiplexed input into output from the BF.

Direct evidence for this assertion was found in the trial-by-trial correlations between firing rate and oscillation amplitude across the power spectrum (Fig. 3E-G). Across trials, the amplitudes of the dominant LFP oscillations at any given task epoch were predictive of the firing rates exhibited by the concurrent population of highly active neurons. Firing rate/LFP power correlations were greatly reduced when trial order was shuffled, suggesting that BF neuron firing rates (i.e., BF outputs) are directly modulated by the LFP oscillatory strengths in different frequency bands (Fig. S3). Effectively, the result evidences an integration of oscillatory input strengths and, further, a transformation of this information into rate-coded spiking output. As shown previously, such BF outputs can drive the responsiveness and patterning of firing in BF target structures such as the cortex in a highly sub-region-specific manner^30^.

The aforementioned evidence for transformation of oscillatory inputs into rate-coded outputs does not preclude a temporal organization, or sequencing, of BF spiking outputs. In fact, combined temporal and rate coding schemes are prominent in the spiking output of other brain systems^31^. We therefore examined whether BF output adheres to a precise sequencing of spiking activity at a finer temporal scale. The analysis focused on BF spiking relative to the theta-frequency component of the BF LFP for several reasons. First, the theta rhythm provides a temporal framework against which beta, gamma, and hi-gamma oscillations are organized (Fig. 1D). Second, theta oscillatory activity was present throughout most epochs of the task (Fig. 3F). Finally, the theta rhythm, in brain regions such as hippocampus, provides a framework for sequenced activity patterns representing trajectories through space^32^,^33^,^34^. Given these findings, we examined the possibility that BF output is sequenced in a similar fashion. Indeed, spiking activity associated with task-epoch-specific increases in firing exhibited phase precession relative to the LFP theta rhythm for a significant population of BF neurons (Fig. 4). To ensure that this result was not epiphenomenal to theta phase resets or shifts at specific task epochs, all BF firing fields that overlapped with any task-epoch bin with a non-uniform theta phase distribution were excluded from the precession analysis (Fig 4C). Critically, phase precession of BF neuron spiking to LFP theta oscillations also generated the expected temporal offsets in cross-correlated spiking activity of neuron pairs whose firing peaks relative to task epoch were offset (Fig. S4). Thus, phase precession yields precise sequencing of BF outputs. In this way, the phenomenon may also support learning processes dependent on the BF^35^ in that such temporal sequencing lies at the core of spike-timing-dependent-plasticity^36^.

**Figure 4.**
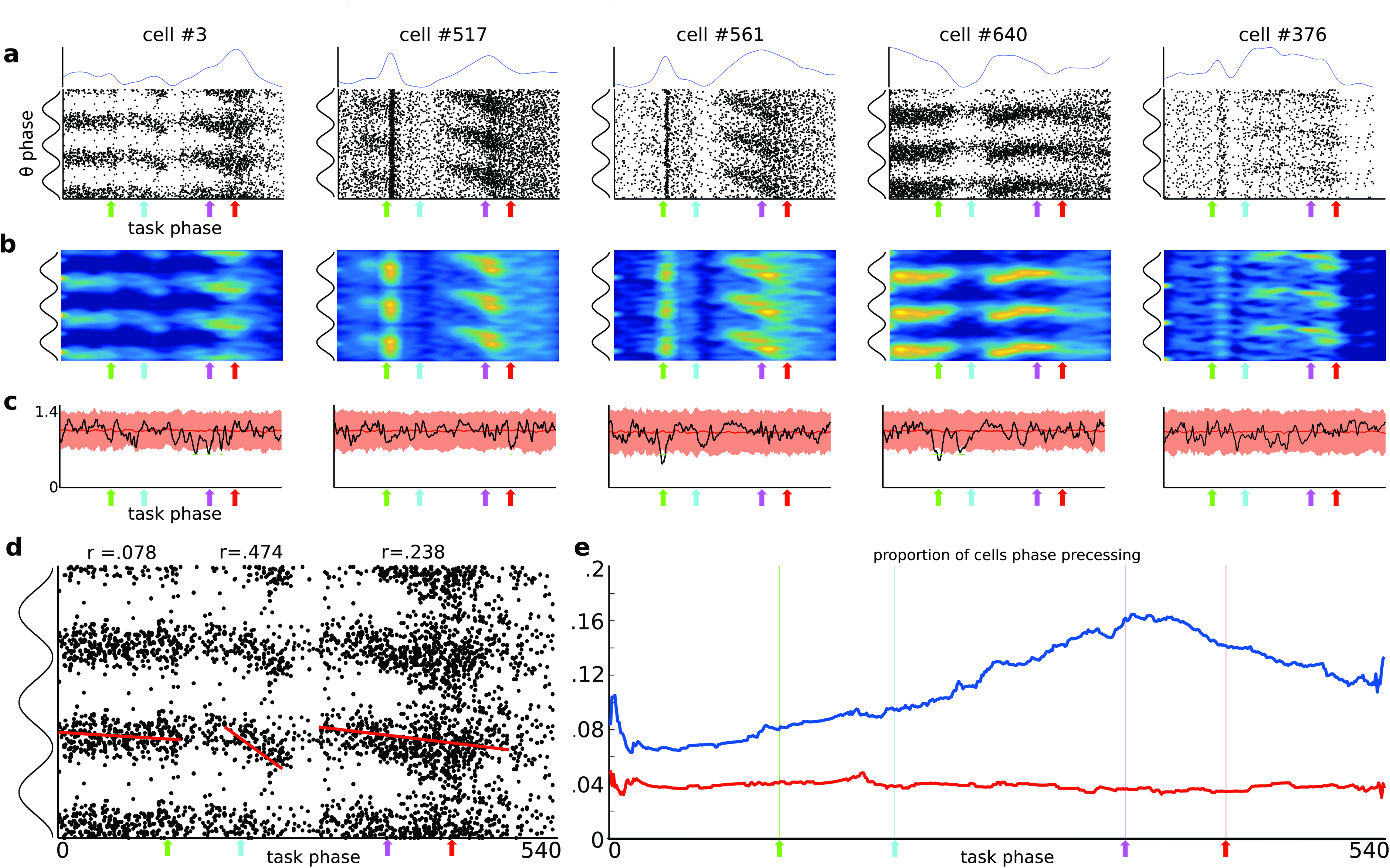
BF neurons theta phase precess across task-epochs. **(A)** Scatter plots, for 5 example cells, of spike times relative to local field potential theta phase (y-axis) and task epoch (x-axis). To aid visualization, theta cycles are repeated three times on the y-axis, and colored arrows depict behavioral events on the x-axis (green/light, cyan/nose-poke, magenta/plate-cross, red/reward). Above the scatter plots, blue lines represent the average firing rate for each neuron across the selective attention task (normalized to maximum rate; y-axis is 0-1). **(B)** Spike density heatmaps of the data given in 3A. **(C)** Theta phases are uniformly distributed across task epochs. Circular standard deviations of theta phases are plotted in black, and those expected by chance in red (10X shuffled, bounded line is a 99% confidence interval). Green dots depict the few task epoch bins with non-uniform theta phase distributions. **(D)** Example cell #3 is depicted again with circular-linear correlations plotted as the slope for each of its firing fields (bins 1-138,185-251, and 291-502). See methods for firing field definition. Note that its second and third firing fields display strong circular/linear correlations between theta phase and task epoch. **(E)** Proportion of neurons that are active (>50% max firing rate) and also exhibiting significant phase precession as a function of task epoch. For each task epoch bin (x-axis), all significant theta precessing firing fields with centers of mass within +/-10 bins were summed (y-axis). Red line indicates the proportion of neurons expected by chance to be precessing at each task epoch.

Notably, phase precession in the BF did not present according to progression through space, as in the hippocampus, nor according to absolute time as has been seen during rapid-eye-movement sleep and salient behavioral events^37^,^38^,^39^ (Fig. 5, Fig. S5). Instead, reliable phase precession was observed when the theta phases of individual spikes were mapped against the concurrent progression of the animal through the task epochs associated with prominent spike rate increases. Thus, phase precession was adaptive to trial-to-trial fluctuations in intervals between major task events such as stimulus onset, choice, and reward obtainment. The result implies that, across trials, the theta phase-specific firing of BF ensembles progresses in step with the progression of the animal through epochs of the task. The task itself requires that the rat fluidly scan the environment, readily apprehend the light stimulus, approach and respond to one of many ports and immediately return to the starting point for a reward. Thus, the ultimate BF output, with its spike sequencing and spike rate magnitudes the products of the integration of multiple oscillatory inputs, follows an abstract chunking process that reflects and/or dictates progression through a behavioral task, rather than the time or space within which the task occurs. Accordingly, BF spiking has been causally linked to the timing of specific behaviors^40^. More generally, similar time-varying progressions through segmented activity states have been seen in cortical^41^,^42^, and subcortical structures^43^,^44^ that integrate multiple sources of information. In this way, processing of information in large scale networks, such as those in which the BF participates, may proceed with some level of independence from the absolute timing of stimuli and actions. Such a level of abstraction may be a critical property for the adaptive flexibility observed in mammalian nervous systems, and the inflexibility observed with BF damage^45^,^46^.

**Figure 5.**
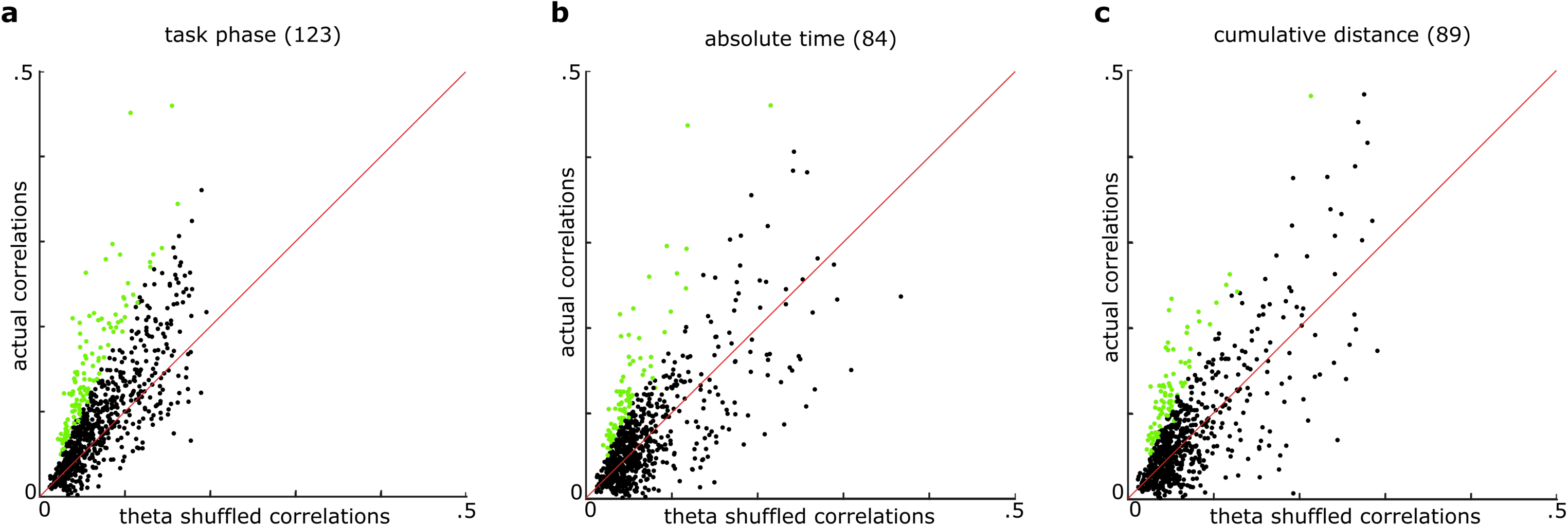
BF phase precession is relative to task epoch progression, not absolute time or spatial location. **(A)** For each neuron the firing field with the maximum circular linear correlation (y-axis) is compared with the average circular linear correlation when theta epochs are randomly shuffled across spike times (mean of 100 iterations; x-axis). Circular/linear correlations are consistently larger for actual task-phase precession than for theta phase shuffled data. Green dots indicate neurons with P-values < .05 for a one sample T-test between actual and shuffled data (123/780 neurons) **(B)** Rather than task epoch bins as in Fig. 5A, absolute time relative to trial onset was used as the linear variable for precession (see Fig. S5 for example neurons). Green dots indicate neurons with P-values < .05 for a one sample T-test between actual and shuffled data (84/780 neurons). **(C)** Rather than task epoch bins as in Fig. 5A, cumulative distance traveled from the trial start was used as the linear variable to test for phase precession (see Fig. S5 for example neurons). Cumulative distance was calculated as the summed Euclidean distance between each position tracking frame from the start of the trial to the frame at which a spike occurred. Green dots indicate neurons with P-values < .05 for a one sample T-test between actual and shuffled data (89/780 neurons).

Together, the foregoing data demonstrate that associative brain regions such as BF employ multiplexing schemes akin to those used in telecommunications to process converging but independent sources of information. Across the population of BF neurons, independently-varying oscillatory afferent inputs are integrated to produce both temporal oscillation in spiking output and to produce rate-coded spiking outputs closely related to the amplitudes of oscillatory inputs. The latter phenomenon provides critical evidence for naturally-occurring transformations between temporal coding and rate coding mechanisms for processing and transfer of neural information. Further, rate-coded spiking output remains temporally organized in the form of specific spike sequences organized through precession of phase-specific firing against the BF LFP’s relatively persistent theta-frequency component.

Across multiple epochs of a selective attention task, this process encompasses highly specific and varying combinations of oscillatory input as well as orthogonal combinations of ensemble spiking output. It is therefore consistent with the implication of BF networks as key to driving the spatiotemporal coordination of cortical activity required to meet a wide range of cognitive demands. The BF, and its cortically projecting constituents, have been known to be vital for normal cognitive function, and its degradation has been linked to neurodegenerative disease, deficits in attention, learning impairments, general dementia, and age-related cognitive decline^47^,^48^,^49^. Thus, it is likely that this complex set of dynamics lies at the core of multiple cognitive processes and its dysfunction may explain why there are such wide-ranging physiological and cognitive impairments associated with damage to this region in humans animals^50^,^51^,^52^.

---

**Figure S1**. Theta, beta, gamma and hi-gamma frequencies are observed across multiple recording days and animals. **(A)** Power spectral density plots are shown for 5,13,12, and 6 recordings in individual animals. **(B)** The average power spectral density of all recording days, with the transparent shading representing two standard deviations. **(C)** Amplitude/amplitude correlations for each animal (color axis 0-1). **(D)** Theta frequency phase/amplitude coupling for each animal. For each frequency (y-axis) 800 millisecond windows of the wavelet transform, aligned relative to a peak in the theta phase, were averaged to get the mean fluctuation in power relative to the peak in theta. Each frequency was then z-scored to show average fluctuations at all frequencies (color axis −3 to 3) **(E)** Randomized phase/amplitude control. The same procedure as in fig. S1D was used, but after the theta-band-filtered signal used to define theta oscillation phases was reversed in time relative to wavelet transforms (color axis −3 to 3). **(F)** Tort’s modulation index (MI) was used to quantify phase/amplitude coupling across recordings. For each frequency band, Tort’s modulation index was calculated for the entire recording. As a control, the LFP phases were flipped in direction relative to the amplitudes, and Tort’s modulation index was recalculated. Scatter plots are the MI scores for each recording and its flipped control. Dashed blue lines represent the median across recordings. **(G)** Summary data for mean MI scores and mean flipped direction MI scores (error bars are +/-1 standard deviation).

**Figure S2**.Average wavelet transform of local field potentials during selective attention task. **(A)** Average wavelet transform for all trials for each animal (5,13,12, and 6 recordings). Green, cyan, magenta, and red arrows demark, respectively, the light flash, nose-poke, plate-cross and stop/reward epochs of the task. All frequencies are z-scored, color axis is from −3 to 3**(B)** Randomized average wavelet transform for each animal. A number of randomly selected segments of LFP, matching the number and duration of the actual trials was used.

**Figure S3**.Fring rate / LFP correlation control. **(A)** Actual mean firing rate / LFP power correlations (color axis: 0-1) as seen in Fig. 2G. **(B)** Firing rate / LFP power correlations when trial order was shuffled (color axis 0-1).

**Figure S4**.Cross-correlogram offsets for simultaneously-recorded neuron pairs correlate with distance between task epoch bins associated with maximal firing. **(A)** Each dot shows the temporal shift to maximal cross correlation (y-axis) for spike times of a pair of simultaneously recorded neurons and the number of time normalized bins (x-axis) between their peak firing rates relative to epochs of the selective attention task. Spike ordering persists despite overlap in the task epoch bins associated with peak task-related firing. The red line indicates the moving median of 20 consecutive task-epoch bins.

**Figure S5**.BF neuron theta phase precession relative to task epoch, time, and space **(A)** For each example neuron (columns), spike rasters were generated relative to theta phase and progression through task phases. **(B)** For each example neuron (columns), spike rasters were generated relative to theta phase and the amount of time passed within each trial. **(C)** For each example neuron (columns), spike rasters were generated relative to theta phase and the cumulative euclidean distance traveled within each trial.

## Supplementary Methods

**Materials and Methods**

**Subjects**

All experimental protocols adhered to AALAC guidelines and were approved by the UCSD Institutional Animal Care and Use Committee and Animal Care Program. Four adult, male Long-Evans rats served as behavioral subjects. Rats were housed individually and kept on a 12 h light/dark cycle. Before experimentation, the animals were habituated to the colony room and handled daily for a period of 1-2 weeks. After this period, animals were placed on food restriction until they reached 85-90% free-fed weight. Water was available continuously. Rats were required to reach a minimum weight of 350 g before surgery and subsequent experimentation.

**Visuospatial Attention Task**

Each day, animals completed 100 trials of a selective attention task in a circular arena with a 1.2 m diameter (Figure 3A). Along the circumference of the arena were 36 light-ports, located at 10-degree intervals and standing 6.5cm above the arena floor. Animals were trained by approximation to remain in a 25 cm circular region in the center of the arena and scan the arena boundary for a light flash (~ 150ms) at a single location. The trial-to-trial probability of a light flash at any given location was defined by a ‘center’ light that is most probable and one of two normal distributions surrounding it (standard deviations 1.25 and 3). Only one distribution was utilized on any given recording day and, across days of testing a single one of these distributions was repeatedly utilized such that several neurophysiological recordings could be obtained under asymptotic levels of performance. Once several recordings were obtained under one distribution, the other distribution was utilized.

Light flash initiation only occurred when the animal was in the center ring and oriented such that the light would fall within a 120-degree space surrounding its longitudinal axis. Thus, the flash location is not always directly in front of the animal, but always within its field of view. Upon detection of a light flash, animals were required to travel to the arena perimeter and to identify the spatial location of the light flash with a nose-poke. Upon returning to the arena center animals were rewarded for correct light source identification with a 1/2-piece Honey-Nut Cheerio (General Mills, Golden Valley, MN). Incorrect identification yielded no reward. Trials associated with failure to travel to the perimeter following light-flash (‘no-gos’) constituted less than 5% of all trials in any animal once asymptotic performance was reached; such trials were not included in the present work’s analyses.

After the animal exhibited correct performance on >70% of trials across several days of training, recording experiments were initiated. At this point, the subject underwent surgery for the implantation of chronic BF and posterior parietal cortex (PPC) single neuron recording wires.

**Surgery**

Rats were implanted with arrays of eight stereotrodes (25 micrometer tungsten with polyimide insulation; California Fine Wire) built into custom-fabricated microdrives. Three such microdrive arrays were implanted in each animal with two targeting left and right BF (AP 0.2 mm, ML 2.8 mm, V 7.0 mm) and one targeting right PPC (AP −4, ML 2.5, V 0.5, 3 animals) or one targeting right BF and two targeting left and right PPC (1 animal). Dorsal-ventral coordinates were chosen to permit slow movement of the recording wires into the desired BF (V 8-9) and PPC (V 0.8-1.5) target areas across days in which the animal was reintroduced to the task. PPC neurons were analyzed previously and are not examined in the context of this paper (Tingley et al., 2014).

**Recordings**

All electrodes were bundled into custom-built microdrives permitting movement in 40µm increments in the dorsal-ventral axis. Signals were amplified at the level of the headstage connection (20X), again at a pre-amp stage (50X), and then to varying degrees, as appropriate, at the amplifier stage (additional 1-15X). Unit signals were bandpass filtered (450 Hz - 8.8 kHz). Candidate spike waveforms (exceeding an amplitude threshold) were recorded using SortClient (Plexon, Dallas, Texas) at a sampling frequency of 40kHz. Waveform discrimination into individual units was carried out manually using Plexons Offline Sorter software.

The animal’s position within the environment was detected from overhead images of the arena at 60 Hz. using Plexons CinePlex Studio. Tracking software picked up light from two differently-colored LEDs clipped to a connector, embedded in the dental acrylic used to fix microdrives to the animal’s skull.

Stereotrode bundles were adjusted across days as necessary to maintain collection of large numbers of high-amplitude action potential waveforms (as many as 60 per day). Data included the present set of analyses were, for all individual animals, associated with different depths (minimum 80 µm separation) to greatly minimize the possibility that single neurons could contribute to the full dataset more than once.

**Behavioral Event Analysis**

Position tracking data was analyzed using a custom Matlab (Mathworks, Natick, MA) guided user interface. Each trial was closely examined to identify the position point associated with initial movement to the light source, and the sharp point of trajectory reversal associated with nose poke. The time points at which the animal crossed back over the perimeter of the center plate and at which the animal stopped to consume reward were determined through semi-automated analysis of positional data using Matlab. Trials in which the animal did not make ballistic, direct runs to and from the site of a nose-poke were not included so that trial-to-trial variability in task epoch durations were kept minimal relative to task epoch mean durations.

**Time-normalization and firing rate calculation**

To enable comparison of neuronal activity across all trials and all behavioral epochs, we utilized time normalization procedure to align neural data for light-onset, nose-poke, center plate return and stop/reward times. Time normalization was accomplished by identifying the average time between light flash to nose-poke, nose-poke to center return, and center return to stop/reward across all trials and animals. On average it took the rodent 0.69s to traverse to the light-port after the light flash. Animals took a mean of 1.41s to return to the plate after nose poke and 0.54s to stop to consume reward after having crossed onto the center plate. We divided these periods into ~80-90ms time bins for each trial. There are slight deviations from these averages for all animals across trials, thus, the bin duration was allowed to fluctuate slightly in order to allow for the behaviorally significant events to consistently occur at the same bin. A 1s period before light-flash and after stop/reward was included in each trial to include stimulus expectation and reward consumption time periods, respectively. By this process, we obtained vectors of time-normalized data in which a pre-light flash period composed bins 1-12, light-flash to nose-poke in bins 13-20, nose-poke to center plate return in bins 21-36, center plate return to stop/reward in bins 37-42, and a post-trial reward period in bins 43-54. For further detail on this time normalization procedure, see Tingley et al., 2014.

**Histology**

Animals were perfused with 4% paraformaldehyde under deep anesthesia. Brains were removed, cut into 50µm sections, and Nissl-stained. The point of deepest electrode penetration was used in conjunction with microdrive adjustment records to determine the range of depths sampled for any given stereotrode bundle placement. All analyses in the present work correspond to histologically identified recording sites in the ventral pallidum and substantia innominate sub-regions of the basal forebrain.

**Amplitude/amplitude modulation**

The local field potential recorded on a single electrode was used for all recordings from each animal. For each frequency band, 1 to 150 Hz in 1 Hz increments, a bandpass Butterworth filter (order = 3) was used with a 2 Hz frequency range (1-3 Hz, 2-4 Hz, 3-5 Hz, etc). These signals were then rectified and smoothed using a moving quadratic mean with a window length two times greater than the maximum frequency, leading to the power envelopes for each frequency band. Correlations were then taken for every pairwise combination of frequency band power envelopes from 1 to 150 Hz, resulting in a 150 by 150 element correlation matrix (Fig. 1C; fig. S1C).

**Phase/Amplitude modulation**

Local field potentials were bandpass filtered for theta frequency (4-9 Hz Butterworth; order = 3) and all peaks in the oscillation were identified. These peaks in theta oscillation were then aligned and averaged, generating the average theta oscillation (grey trace in Fig. 1D). The same alignment of peak locations was then used to average wavelet transforms for frequencies from 20 to 150 Hz. This resulted in an average wavelet transform relative to the peaks in theta oscillations (Fig. 1D).

**Local field potential ‘ratemapping’**

A common procedure with single unit recording is the discretization of recorded spikes to time (or space) bins of a particular size allowing for a mean firing-rate to be calculated for individual bins, also known as rate-mapping. In order to compare local field potential data with single-unit spiking data, we implemented a similar mapping procedure for both data types. For recorded action potentials, a previously described time-normalization procedure was used to calculate firing rates across phases of a behavioral task (Tingley et al. 2014).

In order to compare the local field potential data to this time-normalized spiking data, we first applied a continuous 1-D wavelet transform to the LFP data, allowing for the separation of fluctuations in power across a wide (1-150 Hz) frequency range. Once the wavelet matrix was calculated for each recording, the series of LFP amplitude values occurring between behavioral events for each trial were down sampled by averaging to a predetermined number of time-bins. This allowed for a uniform trial length of 540 bins across all trials for all recordings while preserving the fluctuations in LFP frequency-power across epochs of the behavioral task. To avoid loss of transient high frequency information, the local field potential data was mapped onto 540, instead of 54, bins. All comparisons and correlations between LFP and spiking activity were made after spiking activity was interpolated from 54 to 540 bins.

**Spike/LFP power correlations**

For each neuron, LFP power on each trial (for each frequency band across task epochs) was correlated with the neurons’ coincident firing rates. These correlation values were then averaged across trials (~70 per recording) leading to a 150 (1 Hz increments in frequency; 1-150 Hz) by 1 length vector of average correlation values for each neuron. Each vector (corresponding to one neuron) was normalized by its maximum average value and sorted in the same order as the firing rates in **2E**. Fig. **3G** is a smoothed (average of 30 nearest neighbors; y-axis) heatmap of this matrix (780 neurons by 150 frequency bands). As a control, trial order was shuffled between firing rates and LFP power (Fig. **S3**).

**Phase Precession**

Neuron firing fields were defined as all task epoch bin sequences where a neuron’s firing rate was greater than 50% of that neurons’ maximum firing rate and longer than 10 consecutive bins. For each spike time, the theta phase and task epoch were taken. For each firing field, a circular/linear correlation between theta phase and task epoch (Biostatistical Analysis, J. H. Zar, p. 651) was taken for all spikes that occurred. As a control, the theta phase for each spike time was randomly selected from the distribution of theta phases within each recording, and the same circular/linear correlation was taken (25 iterations). This leads to identical firing fields for each neuron, where the theta phase relationship was effectively randomized. A one-sample t-test (p<.05) was then used to determine significance between the actual circular-linear correlation and the shuffled circular-linear correlations.

To assess how BF neuron phase precession against theta-frequency LFP oscillations relates to the ongoing behavior of the animal, the proportion of active (>50% max firing rate) and significantly precessing (two sample t-test, p<.01) neurons was calculated for each task epoch (Fig. 4E).

For comparison, theta phases for the same spike series were correlated with both absolute time and cumulative distance relative to trial start. Note that the time normalization procedure results in variations in task epoch bin durations across trials such that progression through epochs differs from progression through absolute time. The same subset of action potential times that were used to calculate precession relative to task epoch, were also used for the time and distance calculations. Absolute time is simply the difference in spike time versus trial start time (here, the time of the light flash). For cumulative distance, the summed Euclidean distance traveled from the time of light flash to the time of each spike’s occurrence was determined. Circular/linear correlations between time and theta phase or distance and theta phase were then taken for all spikes within a given firing field

